# Biochemical Barriers on the Path to Ocean Anoxia?

**DOI:** 10.1101/2021.05.17.444596

**Authors:** Stephen Giovannoni, Francis Chan, Edward Davis, Curtis Deutsch, Sarah Wolf

## Abstract

The kinetics of microbial respiration suggest that, if excess organic matter is present, oxygen should fall to nanomolar levels, in the range of the Michaelis-Menten constants (K_m_). Yet even in many biologically productive coastal regions, lowest observed O_2_ concentrations often remain several orders of magnitude higher than respiratory K_m_ values. We propose the *Hypoxic Barrier Hypothesis* (HBH) to explain this apparent discrepancy. The HBH postulates that oxidative enzymes involved in organic matter catabolism are kinetically limited by O_2_ at concentrations far higher than the thresholds for respiration. We found support for the HBH in a meta-analysis of 1137 O_2_ K_m_ values reported in the literature: the median value for terminal respiratory oxidases was 350 nM, but for other oxidase types the median value was 67 μM. The HBH directs our attention to the kinetic properties of an important class of oxygen-dependent reactions that could help explain the trajectories of ocean ecosystems experiencing O_2_ stress.

**IMPORTANCE:** Declining ocean oxygen associated with global warming and climate change is impacting marine ecosystems across scales from microscopic planktonic communities to global fisheries. We report a fundamental dichotomy in the affinity of enzymes for oxygen. The importance of this observation has yet to be fully assessed, but it is predicted to impact the rate at which organic matter is oxidized in hypoxic ecosystems, and the types of organic matter that accumulate. Competition between intracellular enzymes for oxygen may also have impacted microbial strategies of adaptation to suboxia.

## INTRODUCTION

Marine suboxic and anoxic zones are hotspots of microbially-mediated biogeochemical transformations that regulate the nitrogen budget and air-sea fluxes of greenhouse gases of the global ocean (1). Because dissolved oxygen (DO) also organizes the structure and dynamics of ocean food webs, understanding the processes that regulate expansion of suboxic and anoxic zones in response to past and current climate changes is a pressing challenge (2). Suboxic and anoxic zones are embedded within broader oxygen minimum zones (OMZ) that comprise some 8% of the surface area of the ocean. While recent advances in nanomolar-scale DO measurement technologies have enabled precise delineation of the presence of suboxia and anoxia (3), we contend that a perplexing yet fundamental question has been overlooked. Given our canonical understanding of microbial respiration kinetics, why are suboxia and anoxia not a much more pervasive feature of the ocean’s low oxygen zones?

Of biological reactions that consume O_2_, by far the most important, in terms of mass, is carbon respiration. Michaelis-Menten half saturation (K_m_) constants for respiration are typically very low, on the order of a few nanomolar, although higher values have been reported (4) (**Figure 1**). Thus, if labile organic carbon, i.e. compounds that readily can be used as a source of electrons for respiration, is delivered in excess to a microbial ecosystem, DO declines at a rate determined by the respiratory capacity of the microorganisms present and the supply of organic matter. Importantly, the minimum DO attainable should reflect the well-described high-affinity, nanomolar scale K_m_ of microbial respiratory oxidases (5). Other factors that can influence DO in aquatic ecosystems include: photosynthesis, when light is present; oxygen transport by ocean currents and mixing; diffusion, which can limit respiration, particularly in aggregates of cells; impacts of low oxygen on grazing metazoa (6), which require higher oxygen concentrations than bacteria; non-respiratory biochemical reactions that consume oxygen; and abiotic reactions that consume oxygen (7). Nonetheless, DOM formation and oxidation is the mechanistic centerpiece in our fundamental understanding of microbial-scale processes leading to low oxygen states and predictions of global ocean oxygen dynamics.

**Figure 1.**
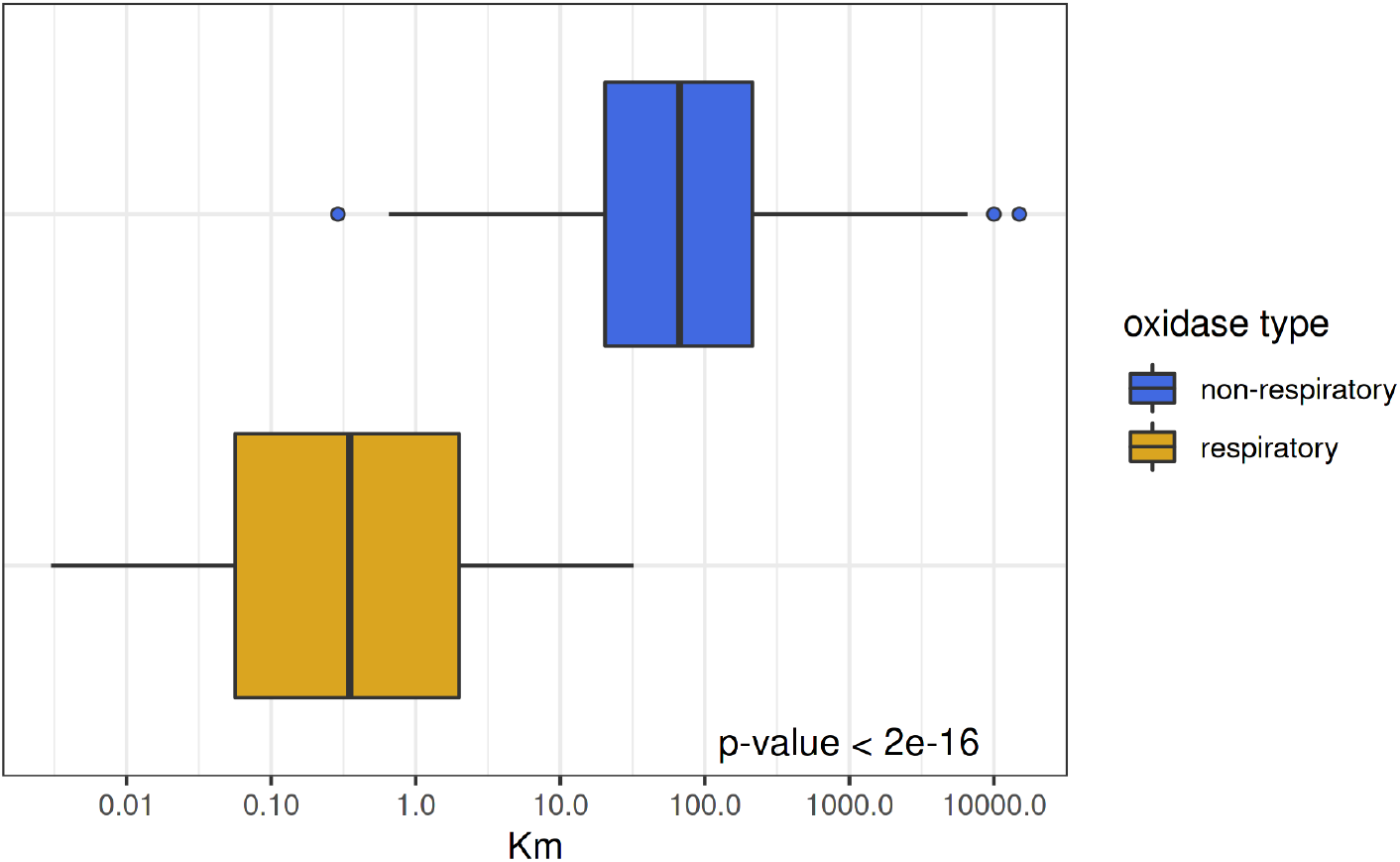
K_m_ values are significantly smaller in respiratory oxidases compared to other oxygenases. K_m_-DO (dissolved oxygen) values for respiratory oxidases (yellow; n=109) and other oxygenases (blue; n=890) are depicted on a log10 scale. The relatively high (e.g. 10^1^ μM) K_m_ values reported for oxidase enzymes indicate a potential bottleneck in the supply of electrons from organic matter to respiration. The reported p-value is from a t-test using the Satterthwaite approximations to degrees of freedom of a linear mixed model fit by maximum likelihood. Data and citations can be found in Table S1.

Eastern boundary upwelling systems (EBUS) represents one of the ocean’s most productive biomes. In these coastal ecosystems, oxygen-poor subsurface waters uplifted from the vertical periphery or core of open ocean OMZs receive elevated organic carbon inputs from surface phytoplankton blooms. **Figure 2** shows relative water volumes for DO concentrations across these systems. Of the ocean’s four major EBUS, only the Peru-Chile Current System in the Eastern Tropical Pacific Ocean persistently exhibits DO-deficient states in continental shelf waters. For both Pacific EBUS, there is an accumulation of water volumes below 100 μM but a sharp drop-off in volume of waters that reach suboxic (<5 μM) or anoxic (~ 0 μM) states in the upper ocean (0-400 m) including continental shelf waters where remineralization and oxygen loss is most active (Fig 2b). The pattern is striking - despite the nanomolar-scale of respiratory K_m_ values, respiration in productive EBUS is able to draw down DO to hypoxic levels but rarely is able to consume the last 10-60 μM DO. While the depth of OMZs extends below 400 m, the failure of suboxic and anoxic volumes to accumulate despite the presence of large volumes of hypoxic water persists when we expand our sampling to 1000 m (Fig 2d).

**Figure 2.**
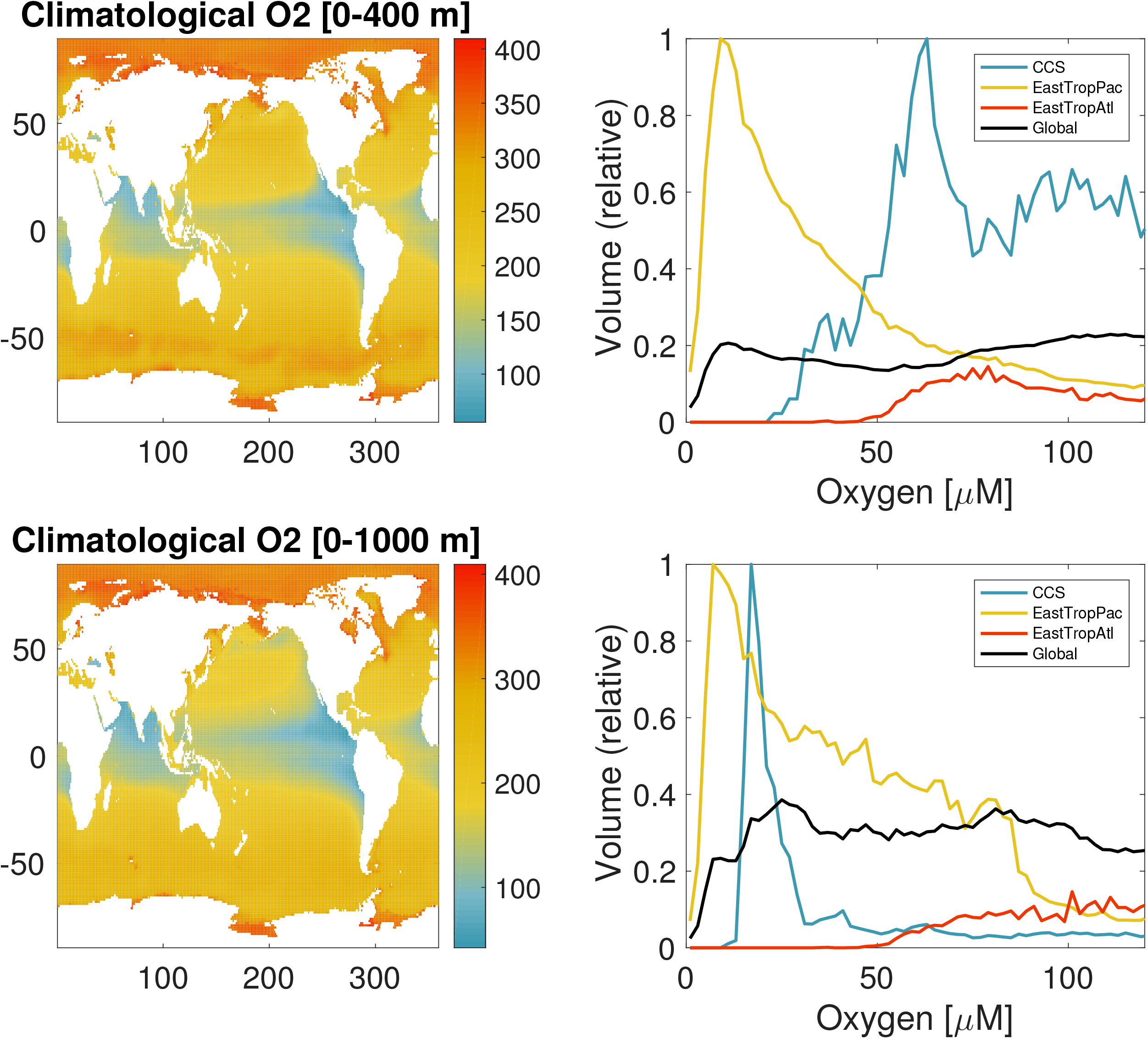
Climatological DO concentrations and distribution of ocean volume across DO concentrations relationships for the California Current System (CCS), Eastern Tropical Pacific (EastTropPac), Eastern Tropical Atlantic (EastTropAtl) and global ocean. (A) 0-400 m, represents generally annual to decadal-scale processes, relative to ventilation and biomass formation by photosynthesis, (B) 0-1000 m, encompasses decadal to centennial-scale processes.

To explain the observations in **Figure 2**, we propose the ***Hypoxic Barrier Hypothesis (HBH)*, which states: *dissolved O_2_ kinetically limits the activity of oxygenase enzymes involved in the breakdown of organic matter, in the range of oxygenase K_m_ values (median value 67 μM) causing a decline in DOM oxidation rates in ecosystems experiencing oxygen stress, and an accumulation of DOM that is catabolized by pathways that require oxygenases*.** The *HBH* ascribes the decline in O_2_ frequency distributions of suboxic and anoxic waters (Figures 2 and S2) to fundamental biochemical properties of cells, particularly the mechanisms by which oxidative enzymes cleave semi-labile organic matter, making it accessible to further oxidation.

### Oxygen depletion by respiration in aquatic systems

How far can respiring marine bacteria lower oxygen concentrations when they are provided with an ample supply of reductant for respiration, as would be expected for plankton in the presence of an excess of labile organic carbon? The Pasteur Point is an influential concept based on the observation that facultative anaerobes switch to fermentation at ca. 2.2 μM O_2_, approximately an order of magnitude below the average K_m_ for oxygenases (Fig 1), steep declines in suboxic and anoxic water volumes in EBUS (Fig 2), and the ca. 25 μM inflection in cumulative frequency distribution of DO observations recorded in CCS (Fig. S2). Newer information suggests that the limits of bacterial respiration are in the nanomolar range. This is consistent with the observation of high affinity cytochromes that exhibit O_2_ K_m_ between 3 and 200 nM (8; Figure 1). Higher O_2_ Michaelis constants have been sometimes been reported for marine bacteria, but it has been suggested that higher values obtained with whole cells reflect diffusion limitation, which can be expected to inflate apparent O_2_ Michaelis constants in proportion to cell sizes and respiration rates. Stolper et al. showed that *E. coli* cells could grow at less than 3 nM O_2_, a sufficiently low concentration to limit growth by diffusion, but high enough to sustain growth through O_2_ respiration (8). Our meta-analysis of indicates that cells grown on highly labile carbon compounds such as glucose display whole cell O_2_ K_m_ values extends well into nanomolar O_2_ concentrations. We conclude that a substantial background of observations and theory support the conclusion that the respiration rate of chemoheterotrophic cells should not by limited by O_2_ at concentrations found in ocean hypoxic zones or the Pasteur Point at ca. 2.2 μM.

The accumulation of hypoxic and scarcity of suboxic or anoxic volumes in the ocean nonetheless suggest that negative feedbacks between oxygen decline and respiration may be at play. Direct measurements of microbial O_2_ K_m_ in natural systems are rare but available evidence point to K_m_ values far higher than the nanomolar values reported from laboratory cultures with labile carbon sources. Working in the Arabian Sea OMZ, Keil et al. (9) observed an apparent K_m_ of 20 μM O_2_ for microbial community respiration. In the Nambian and Peruvian OMZ, Kalvelage et al. (10), reported a linear decline in respiration rate between 20 and 0 μM O_2_. In Chesapeake Bay, a hypoxia-prone system, microbial respiration rates saturate at [O_2_] above 25 μM (11), a pattern that we have similarly found for the CCS OMZ (Figure S3). Holtappels et al. (12) further reported linear declines in respiration rates between 14 and 1 μM O_2_ in waters collected from a fjord in Denmark. These results are surprising because researchers using the same methods have also found many instances of nM K_m_ values for microbial respiration. This suggests a bimodal distribution of O_2_ K_m_ values that differ by upwards of three orders of magnitude. Telescoping out further, global models of ocean O_2_ and carbon export converge on K_m_ values of between 4 and 20 μM O_2_ in order to optimize fit between model and observations (13, 14). What accounts for the disparity between accumulation of hypoxic water volumes, the μM scale K_m_’s reported from natural systems and used to fit models, and nM scale K_m_’s predicted by respiratory oxidases?

Biochemistry offers a mechanistic explanation for this apparent disparity. There is evidence in the scientific literature suggesting that microbial respiration of some types of organic matter is slows when oxygen concentrations fall low enough to inhibit catabolic oxygenase enzymes. Kroonman et al. (15), studying 3-chlorobenzoate degradation by the bacterium *Alcaligenes*, reported two K_m_ values for O_2_ uptake. They attributed the lower value (65 nM) to respiration and the higher value (7-17 μM) to the activity of dioxygenases. Leahy and Olsen (16), studying toluene degradation by *Pseudomonads*, also reported biphasic kinetics for toluene catabolism as a function of oxygen concentration. The slope of the oxygen response declined with an inflection at 20 - 30 μM O_2_. In both of these cases the behavior of the cultured cells oxidizing recalcitrant compounds is remarkably similar to the generalized behavior of ocean ecosystems approaching hypoxia.

To further explore the distribution of O_2_ K_m_ values among biological reactions, we conducted a metanalysis of published data, shown in **Figure 1; Figure S1A; Table S1**. For O_2_ K_m_ values reported in the literature, the median value for terminal respiratory oxidases was 350 nM, but for other oxidase types the median value was 67 μM. The difference of ~100 fold in median values was supported by a p-value of <2e-16 in a t-test of the linear mixed-effect model coefficient comparing the log-transformed O_2_ K_m_ values (17). The bimodal distribution of O_2_ K_m_ observed at the enzyme scale is also repeated in whole cell studies. Cells that are grown on more complex organic carbon sources have a median respiratory K_m_ value of 20 μM, while cells grown on highly labile organic carbon such as glucose have median K_m_ of 690 nM (Figure S1B).

Many enzymes that catalyze the biological breakdown of organic matter use oxygen as a substrate, yielding partially oxidized products that are metabolized further through catabolic pathways. These enzymes are often classified as either monooxygenases (mixed function oxidases) or dioxygenases. Enzymes in both families evolved to use O_2_ as a substrate, but monooxygenases incorporate a single oxygen atom into the substrate, reducing the second atom to water, whereas dioxygenases typically add both atoms of the reacting O_2_ to the product. A geochemically important example of a monooxygenase is the heme-dependent Mn peroxidase that catalyzes oxidation of lignin, a phenolic oligomer. Fungal ligninases belong in the heme-dependent peroxidase superfamily (18). Ligninases evolved in the Paleozoic, and it has been postulated that their origin resulted in widespread biodegradation of wood, causing the end of the carboniferous period, and rises in global atmospheric CO_2_ (19). Another superfamily of oxidases, the flavin-dependent monooxygenases, are among the most diverse and prevalent proteins known. They catalyze a wide range of reactions, for example hydroxylation, Baeyer–Villiger oxidation, oxidative decarboxylation, epoxidation, desulfurization, sulfoxidation and oxidative denitration (20). Many of the reactions catalyzed by flavin-dependent monooxygenases initiate the catabolism of compounds that are otherwise recalcitrant to oxidation. These enzymes share a common mechanism in which reduced flavin reacts with oxygen to produce a flavin C4a-(hydro)peroxide that then reacts with electrophilic or nucleophilic substrates, typically resulting in the consumption of one diatomic oxygen molecule, the addition of an oxygen atom to the substrate, and the release of water. Also important are the dioxygenases, which belong to a different protein family also feature prominently in DOM degradation particularly for aromatic compounds. These protein families came to the attention of oceanographers recently when it was discovered that cells of one of the important oceanic bacterial clades, SAR202, harbor expanded clusters of paralogous genes from both of these protein types (21). It has been proposed that these enzymes participate in the oxidation of semi-labile organic matter, initiating its breakdown.

The meta-analysis of O_2_ K_m_ values presented in Figure 1 suggests that micromolar DO sensitivity is ubiquitous across metabolic processes. Among the “other oxidase” types, we observed no clear trends that associated O_2_ K_m_ values with protein families sorted by COGs or other precise functional groups, such as enzyme commission classifications. In contrast, cytochrome respiratory proteins with heme cofactors consistently displayed a much higher affinity for oxygen that other protein types. Phylogenetically, the distribution of O_2_ K_m_ values included diverse bacteria, including Proteobacteria, Actinobacteria, Firmicutes, and Cyanobacteria, as well as eukaryotic organisms including fungi, humans, and other chordates. O_2_ K_m_ values showed a 100-fold difference between respiratory and non-respiratory oxidases regardless of taxonomic group. We conclude that this pattern is robust to phylogenetic bias in O_2_ K_m_ value sampling. While our focus in the current work is marine systems, the HBH in principle applies to all ecosystems.

### The role of oxidases in organic matter degradation

Oceanographers classify organic matter by its half-life, frequently using the category “labile dissolved organic matter” (LDOM) to describe dissolved organic matter that is oxidized in minutes to hours, or at most a few days, while the term “semi-labile” (SLDOM), and sometime “recalcitrant” are used to refer to dissolved organic matter that persists longer, but is eventually oxidized. Here we introduce a new term, “oxygen-dependent DOM” (ODDOM) to describe DOM that is catabolized via reactions that require the activity of oxygenases and thus are susceptible to inhibition when O_2_ concentrations reach values in the range of ca. 10-100 μM. We’ll confine the discussion to dissolved forms of organic matter, although most organic matter enters ecosystems as particulate organic matter (POM) and is subsequently converted to DOM before being used by microorganisms. Implicit in the above categories is the idea that different kinds of organic matter are accessible to biological oxidation through different mechanisms and at different rates.

The HBH is consistent with the distribution of ocean anoxic zones if one assumes organic matters supplies are uneven. If LDOM is oversupplied relative to oxygen, for example by high rates of export production in systems with restricted circulation, then the activities of respiratory terminal cytochrome complexes would be expected to readily draw down DO to nM concentrations in accordance with their nM K_m_ values. In natural systems, LDOM are rapidly depleted. As hypothesized, the activities of non-respiratory oxidases limit the supply of reductant to respiratory oxidases. This acts as a bottleneck that slows the rate of respiration as DO declines. With sufficient time, DO should reach minimum values as expected from nM K_m_ values of respiratory oxidases. Such conditions can be met in the core of OMZs that have been isolated from the atmosphere over decadal to century time scales, and evidence of this can be seen in Fig. 2.

The large disparity in K_m_’s we report between respiratory oxygenases and other oxygenase types has implications for microbial cell evolution and metabolic regulation at the cellular level. Inside of cells respiratory oxygenases could outcompete other oxygenases, exacerbating the slowing of some oxygen-dependent cellular processes at low oxygen. To avoid this, cells may have evolved metabolic regulation that avoids such competitive interactions, for example by shifting to alternate electron acceptors before O_2_ is depleted (22). This topic, which needs exploration, could help us understand how microbial cells have adapted to suboxic environments, which are far more common in the ocean than anoxic environments.

### Testing the HBH

The HBH sets forth a number of central predictions that are testable by experimentation, observation, and modeling. The impact of biphasic oxygen dependence predicted by the HBH should be manifested as a broad potential for oxygen to limit microbial respiration across hypoxic systems in the range of oxygenase K_m_ values (median value 67 μM), when LDOM is depleted, but not if excess LDOM is present. To test that prediction, we measured rates of respiration (oxygen uptake) in water samples from the Northern California Current System OMZ, where DO minimum reach only ~5 μM, well above canonical nM K_m_ for cytochrome oxidases. DO was increased by the simple expedient of allowing air to be momentarily entrained during filling (**Figure S3**). In our experiments, and other similar experiments we found among published work, the addition of DO caused respiration rates to rise relative to controls. This observation could be attributed to the limitation of respiration by diffusion (20), but alternatively, it could result from mechanisms described in the HBH model we propose.

There are many other experimental avenues to testing the HBH that have not been explored. **Figure S2** scratches the surface of what could done with field experiments and mesocosms to verify predictions of the HBH. For example, experiments that test the biological availability of DOM at high (e.g. 200 μM) and moderate (e.g. 20 μM) DO could challenge these ideas. Mechanisms invoked by the HBH would lead to changes in the chemical composition of DOM as DO declines: the ratio of LDOM to ODDOM should decrease as DO approaches the K_m_ values of catabolic oxygenase enzymes for O_2_. Measurements of DOM chemistry could determine whether these changes occur as predicted. ODDOM, a term coined herein to segregate DOM into categories by chemical composition and oxygenase involvement in catabolism, is at present a theoretical concept, albeit grounded in the fundamentals of biochemistry. Although chemical oceanographers do not at present measure ODDOM, in principle methods such as high resolution NMR, HPLC, and LC-MS/MS could be applied for this purpose, and could be used to test predictions of the HBH. Omics approaches, including functional genomics, provide an avenue that could be applied in marine systems to measure the expression and activity of oxygenase enzymes involved in ODDOM metabolism, and to characterize of the responses of plankton cells and communities to suboxia.

The HBH has broader implications that could be explored with global data. It posits that rates of oxygen loss and DOM oxidation slow as DO approaches hypoxia, setting the upper bounds for the size of oceanic anoxic zones and organic carbon pools within. This can be evaluated in detail by modeling studies that test the sensitivity of model-data comparisons to changes in assumptions about microbial kinetic constants for oxygen.

### Alternatives to the HBH

While we propose HBH to explain declines in oxygen frequency distributions at unexpected high values (Figs. 2 and S2) and the rarity of suboxia and anoxia across productive, low oxygen EBUS, alternate scenarios could explain this phenomenon. For example, consistent barriers to oxygen diffusion to the terminal oxidases of respiratory systems, or feedback mechanisms involving the production of sulfides and/or depletion of DO in microhabitats, or oxygen limitation of metazoan grazing, could play a role in suppressing respiration at low oxygen concentrations. Alternatively, there may be constraints on supply of organic carbon or positive feedbacks on the resupply of DO by advection or diffusion as DO approaches hypoxia.

### Public interest and policy

The relevance of this issue to public interests in ecosystem management could not be more profound. Ocean deoxygenation, the decline in ocean oxygen inventories, has emerged as a leading pathway for climate change impacts in the sea. This decline has been linked with expansion of hypoxic and anoxic zones. Oxygen deficient zones are hotspots of biogeochemical transformations whose growth can have profound impacts on marine biodiversity, vertical organic carbon flux, the sustainability of fisheries and feedbacks that govern ocean nitrogen budgets and flux of radiatively active N_2_O. The ability to accurately forecast such ecosystem changes is central for informing responsive climate change mitigation and adaptation policies. However, the disagreement between observations and the textbook understanding of microbial respiration raises fundamental questions about the mechanisms that underlie our conceptual and numerical models of the ocean dynamics as climate change intensifies. The HBH offers a testable framework for examining a potentially flawed fundamental principle that governs our thinking about OMZ formation. If this hypothesis is correct, it will open previously overlooked avenues of research at the intersection of oxygenase enzyme evolution, oxygenase-dependent metabolism in microbial communities, and OMZ dynamics.

### Conclusion

If these ideas have the power to even partially explain the kinetics of ocean oxygen depletion, they could contribute to a better understanding of climate change impacts on ocean deoxygenation and DOM chemistry. The data in Fig. 1 show us that the HBH is founded on sound basic principles, but the impact of oxygenase “barrier” we describe is relative to many other processes, mentioned above, that can also slow respiration, most notably diffusion. Sorting out the magnitude of catabolic oxygenase enzyme contributions to DOM oxidation, whether that number be large or small, will help us assess how the trajectories of aquatic systems experiencing oxygen declines are shaped by the fundamental biochemistry described in the HBH.

## MATERIAL AND METHODS

### Data collection

Scientific literature was mined for characterized oxygenase enzymes with published K_m_ values for dissolved oxygen for both individual enzyme assays as well as whole cell assays (Table S1). Metadata, including enzyme name and host scientific organism name, were extracted from each article. We used a combination of BRENDA enzyme database, uniprot protein database, and KEGG database searches to determine putative protein accessions, KEGG ortholog IDs, and EC numbers associated with the published enzyme data. Repeated entries for the same organism-protein pairs were included due to the various testing conditions per study.

Basin and global inventories of DO volumes were compiled from the World Ocean Atlas 2018 (https://www.ncei.noaa.gov/products/world-ocean-atlas. Dissolved oxygen observations (5-400m) from CTD profiles were compiled from Chan et al. 2008, https://www3.mbari.org/bog/, and https://www.calcofi.org/ for the northern (n=107,032 1950 to 2006), central (n=4,372 1997-2013), and southern (n=4,372 1997-2013) CCS, respectively.

### Respiration rate experiment

Water samples were drawn from above and within the CCS OMZ (46 47.56°N, 125 11.83°W, 1000m station depth) and filled into 300ml borosilicate glass BOD bottles that each contained an oxygen optode dot (PreSens Precision Sensing GmbH). On filling, DO in a subset of samples initial [DO] were increased by allowing air to be entrained momentarily in the Niskin outflow tubing. Bottles were incubated in a ~6°C water bath in the dark. DO change over 48 was measured through the glass via detection of phase shift luminescence.

### Boxplot generation

Km-DO values were split into two primary groups dependent upon general protein function, either respiratory oxidases or non-respiratory oxidases. K_m_-DO values were also gathered for whole cells, whereby labile and semi-label carbon sources were compared, mirroring the respiratory and non-respiratory individual enzyme assays. Plots were generated with the grouped K_m_-DO values using R v4.0.2 (23) and the ggplot2 (24). A linear mixed effects model was used to control for repeated O_2_ K_m_ measurements from the same organism, with the formula log(K_m_) ~ oxidase type + (1 | organism). The model was fit using maximum likelihood, and the t-test to confirm significant difference of the coefficient for oxidase type was done using Satterthwaite’s method for degrees of freedom (17). Log (natural) transformed values were used to approximate normality in the data. R code for this analysis can be found at github repo: https://github.com/davised/HBH-2021.

## ACKNOWLEDGEMENTS

We thank the reviewers, John Coates and Dave Valentine, for their many useful comments. John Coates offered the important insight that intracellular competition with respiratory oxygenases could contribute to the inhibition of non-respiratory oxygenases. This work was funded by the National Science Foundation grant DEB-1639033, NOAA grant NA18NOS4780169, a SciRIS award from the Oregon State University College of Science, and a grant from Simons Foundation International. No authors declare any real or perceived financial conflicts of interests.

## SUPPLEMENTAL MATERIALS

**Table S1.** Literature reports of K_m_ values for oxygenase enzymes and terminal respiratory oxidases used to construct Figs 1 and S1.

**Fig. S1.** Oxidase density plot. This plot illustrates the distribution of oxygenase K_m_ values reported in the literature for respiratory and non-respiratory oxidases (A) compared to whole cell assays including labile and semi-labile carbon sources (B).

**Fig. S2.** Cumulative frequency distribution of DO observations from continental shelf depths (5-400 m, i.e. above the OMZ) for the northern 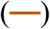, central 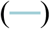, and southern 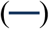 California Current System (CCS). Note the rarity of DO observations < 25 μM and an apparent hinge in the frequency of observations as DO increases beyond that concentration.

**Fig. S3.** Examples of respiration rates from field collected samples that exhibited non-saturating dynamics at 10’s of uM [O_2_]. In each instance, respiration rates were assayed in samples where [O_2_] was manipulated independently. Data from the CCS were measured via O_2_ optode equipped glass bottles from water samples collected within the OMZ. Oxygen was increased by allowing air to be momentarily entrained as bottles were filled.

